# Adiposity in Adolescence has Detrimental Effects on Developing Brain Structures and Organization, Communication, and Controllability of Resting-State Networks

**DOI:** 10.64898/2026.06.22.733806

**Authors:** Matthew Risner, Emma Martin, Catherine Stamoulis

## Abstract

Adiposity in adolescence can have detrimental effects on neural maturation, and is associated with incompletely understood alterations in brain structures and circuits. To address this important knowledge gap, N=3,341 youth from the Adolescent Brain Cognitive Development (ABCD) two-year follow-up cohort (median(IQR) age=12.0(1.1) years; 51.0% females), with structural MRI and resting-state fMRI were studied. Topological properties reflecting network strength, resilience, efficiency and modularity, information transfer, regional feedback costs associated with controllability of network dynamics, and morphometric characteristics were examined as a function of body mass index (BMI), body roundness index (BRI), overweight, obesity, and persistent excess BMI (across assessments). Youth who slept less, spent more time on electronic devices, were Hispanic and/or from lower-income families had higher BMI/BRI (β=0.07-4.95, 95%CI=[0.03,7.05]), and higher odds of overweight or obesity (aOR=1.01-1.72, 95%CI=[1.01,2.28]). Higher BMI/BRI, overweight, obesity and/or persistent excess BMI were associated with weaker and less resilient networks supporting decision-making, control, emotional regulation, reward processing and social function (β=-0.11 to −0.01, CI=[−0.16, −0.01]), and more topologically fragile thalamus and basal ganglia (β=0.05-0.10, CI=[0.01,0.14], p<0.03). They were also associated with lower control costs in cognitive and topological hubs (including the precuneus and prefrontal regions) that play central roles in regulating brain dynamics, aberrant information transfer in limbic, frontoparietal, salience, somatomotor and cerebellar regions, and lower thickness and/or volume of distributed (including prefrontal) regions, and functional hubs (β=-0.14 to −0.02, CI=[−0.18,-0.01]). Thus, adiposity in adolescence is associated with widespread structural and alterations of brain networks supporting developing cognitive processes, and fundamental mechanisms that control these networks’ dynamics.

## 1. INTRODUCTION

Excess weight affects one in six children in the US, and its prevalence increases with age [1]. Currently, over 20% of adolescents have obesity, a highly alarming statistic with profound negative implications for the individual and society [2]. Although genetic factors increase predisposition, environmental risk factors, poor diet, insufficient sleep, lack of physical activity and increased screen time are significant drivers of the obesity epidemic [3]. In addition to widespread detrimental effects on physical health, excess weight has also been linked to cognitive deficits and mental health issues in youth, through mechanisms of action on the developing brain that are incompletely understood. This gap in mechanistic knowledge is a significant barrier to the development of science-based interventions that are tailored to the individual, as well as necessary health, environmental and regulatory policy changes that are critically needed to combat the obesity epidemic - a growing global issue [4–5]. This study aimed to address this gap, and investigated the neural correlates of excess weight across domains, including brain structure, circuit organization and communication (information transfer). It also used a powerful modeling approach to elucidate the impact of excess weight on unobserved regulatory processes that control brain function. The study focused on adolescence, a sensitive period of brain development during which exogenous and endogenous factors, such as excess weight, increase the risk of circuit miswiring, cognitive deficits, and mental health issues.

Adolescence is a vulnerable period for mental health, and common disorders often emerge in adolescence, likely as a result of aberrant neurobiological changes, including disrupted or delayed neural maturation and miswiring of brain circuits that play critical roles in mental health [6]. Prior studies have linked excess weight to widespread mental health problems in adolescents, including higher incidence of depressive symptoms, increased risk for anxiety disorders, and suicidality [7–18]. Although aberrant circuit alterations and structural changes in youth with excess weight are incompletely understood, these studies point to complex relationships between adiposity, neuroinflammation and aberrant brain changes underlying mental health problems [19–24].

Excess weight has also been linked to cognitive deficits, especially in domains that continue to evolve in adolescence. Youth with overweight or obesity may have lower inhibitory control and cognitive flexibility, higher emotion-driven impulsivity, deficits in executive function, impaired emotion and reward processing and regulation, and lower overall academic achievement [25–32]. Prior studies have shown that these associations are partly driven by the adverse effects of excess weight on the developing brain, including structural brain alterations (such as abnormally lower cortical thickness and/or volume, and disrupted white matter integrity [33–36]), and aberrant topological differences in functional networks, especially those that undergo heightened maturation in adolescence [37–45]. Most studies have examined effects of excess weight on the strength of network connections, while far fewer have examined other topological properties. The relatively few studies that have examined these networks’ topological patterns and organizational properties have found topological alterations in network modularity and efficiency that have detrimental implications for cognitive processing across domains [42–45].

The majority of prior investigations of brain-BMI relationships in adolescents have been on relatively small samples, but their findings have been recently been replicated in large adolescence samples, including that of the Adolescent Brain Cognitive Development (ABCD) study [46]. This historically large study has also provided new knowledge on relationships between excess weight and brain development. Studies have reported negative associations between BMI and executive function that are partially mediated by lower thickness of the prefrontal cortex [35,47], and reduced gray matter volume in youth with obesity [48], including in regions that support working memory [49]. Another study showed that the microstructure of the nucleus accumbens is associated with obesity and predicts future weight gain [50–51]. BMI has also been associated with stronger connections between the putamen and the salience network, but this relationship dependent on youth race and their parents’ marital status [52].

Overweight and obesity have also been linked to brain alterations that are partially mediated by sleep disordered breathing [53]. Studies focusing on sociodemographic factors have reported amplified adverse effects of obesity on the brain in youth from low-income families [54–55], and associations between neighborhood deprivation and BMI that are mediated by structural brain alterations [56]. Our previous study on the baseline (pre/each adolescent) cohort of the ABCD has shown that both overweight and obesity are associated with topological and morphological brain alterations [45], in agreement with prior ABCD-based studies on structural correlates of adiposity [57], and that physical activity has positive effects on brain regions that are adversely impacted by excess weight [58].

While our prior work examined pre/early adolescents, the present study focused on the 2-year follow-up cohort of the ABCD, and thus youth in mid puberty, a sensitive period of accelerated maturation of circuits that play key roles in reward processing (including food reward), emotional regulation and decision-making (especially the prefrontal cortex) undergo substantial reorganization [32, 59–62] . First, we hypothesized that negative associations between BMI (and body roundness) structural brain features, and aberrant topological alterations network remain consistent as youth age, and are amplified in youth with persistent excess BMI in pre/early and mid puberty. We further hypothesized that excess BMI also affects the developing brain’s intrinsic (task-independent) dynamics, which play a ubiquitous role in cognitive function [63–65], and are controlled by endogenous biochemical processes that may not be directly measurable in real time in the intact human brain. In a recent study on both the baseline and 2-year follow-up ABCD cohorts, we used a novel to Neuroscience closed-loop control framework to model these latent processes and elucidate the adolescent brain’s controllability [66]. We showed that the adolescent brain’s dynamics are controlled by a sparse, and potentially developmentally invariant set of topological and cognitive hubs that include both developing and developed regions. Here, we hypothesized that excess weight adversely impacts latent (biochemical) processes that play key modulatory roles in neural dynamics, and its effects are reflected in alterations of the brain’s controllability. Finally, we hypothesized that, through its detrimental effects on network organization, excess weight also impairs brain communication, a fundamental mechanism of brain function and information processing, and investigated BMI-related alterations in information transfer (effective connectivity and flow) between brain regions. The overarching goal of this study was to gain mechanistic (and comprehensive) knowledge on detrimental impacts of excess weight across dimensions of adolescent brain function, from structural hallmark characteristics to fundamental mechanisms underlying the organization, dynamics and communication of developing brain circuits, including controllability. To the best of our knowledge this is the first investigation to examine and integrate these brain dimensions in the context of excess weight in adolescence.

## 1. METHODS

Neuroimaging, anthropometric, and demographic data from the 2-year follow-up cohort of the ABCD study were analyzed (all data were from release 5.0; at the time of this study’s analyses release 6.0 data had not yet been shared with the community). These data are publicly accessible through the National Institute of Mental Health Data Archive (NDA). The study was approved by the Institutional Review Board.

### 2.1 Participants

Participants with underweight (≤5th percentile), clinical MRI abnormalities (identified by the ABCD study as abnormalities requiring further evaluation), and/or a diagnosis of a neurodevelopmental or psychiatric disorder were excluded [39]. Furthermore, youth with biologically implausible BMI estimates due to errors in weight and height (defined by the CDC as BMI z-scores <-4 or >+5) and/or negative changes in height from baseline to follow-up were excluded. Finally, participants with poor quality imaging data (based on the quality control criteria set by the ABCD study and conservative quality criteria in our pipeline, including the criterion of at least one 5-min fMRI run with ≤10% frames censored for motion) were also excluded. Following these exclusions, a final analytic sample of N = 3,341 adolescents was identified (median (IQR) age at fMRI scan = 12.0 (1.1) years; 1,705 (51.0%) girls). This sample included n = 1,953 youth (58.4%) who also had baseline data that met all criteria for inclusion.

### 2.2 Measures of Obesity and Excess BMI

BMI was calculated from weight and height and was standardized for age and sex as z-scores. Percent median BMI (%mBMI) was estimated as a measure of each participant’s BMI relative to the population. Body Roundness Index (BRI) was also analyzed, and was calculated using the equation [67]: BRI = 364.2 – 365.5*√(1- (waist circumference(cm)/2*π)^2^/0.5*height(cm)^2^.

In addition to these continuous measures, youth were classified as having normal BMI, overweight or obesity, using cutoffs estimated separately for boys and girls, based on age- and sex-specific growth curves [2]. These groups were represented by binary variables (0 = normal weight, 1 = overweight or obesity). A third binary variable was used to contrast obesity and overweight (0 = overweight, 1 = obese). Finally, a binary variable contrasted persistent excess BMI over time to persistent normal BMI, i.e., 1 = excess BMI both at baseline and year 2 follow-up, vs 0 = normal BMI at both assessments.

### 2.3 Neuroimaging data

Structural MRI (T1- and T2-weighted) and resting-state fMRI were collected at 21 ABCD sites using 3.0T GE, Siemens, or Philips scanners (fMRI repetition time (TR) = 800 ms; resolution = 2.4 mm isotropic). Each participant completed up to four 5-minute resting-state fMRI (rs-fMRI) runs. Minimally preprocessed data by the ABCD Data Analysis, Informatics & Resource Center [68] were further processed in our custom pipeline [69]. All analyses were conducted using the software Matlab (release R2025a, Mathworks, Inc).

#### 2.3.1 Resting-state fMRI data and topological property estimation

Neuroimaging data were processed using the Next Generation Neural Data Analysis (NGNDA) pipeline [69]. Each fMRI was coregistered to the participant’s T1w MRI, normalized in MN152 space, corrected for motion, denoised, and downsampled to 1088 parcels. Three atlases were used for this purpose: the Schaefer cortical atlas (1000 parcels) [70], Melbourne subcortical atlas (54 parcels) [71], and a probabilistic cerebellar atlas (34 parcels) [72]. All data were harmonized through signal normalization to account for scanner differences and resulting BOLD signal amplitude differences.

Frames were censored for motion, using a displacement threshold of 0.3 mm. Only runs with ≤10% of censored frames were analyzed [39]. Each participant’s best run was selected based on lowest median connectivity, typically coinciding with the run with the lowest percent of frames censored for motion (median (IQR) = 1.3 (4.0)% in this sample). For participants with multiple usable runs (n = 2,580; ∼76% of the sample), a second run was analyzed (median (IQR) percent of frames censored for motion = 1.3 (3.7)%), for reproducibility purposes, and thus a total of 10 min resting-state activity.

Pairwise peak cross-correlation between two signals was used as a measure of functional connection and was calculated for all pairs of parcel signals, resulting in 1088X1088 fully connected graphs, which were then thresholded to eliminate spurious connections. For this purpose, a conservative, brain-wide threshold corresponding to the moderate peak cross-correlation outlying value (equal to median + 1.5*IQR) [58] was used. Based on this threshold, <20% of resting-state connections were typically retained. Topological properties at the whole-brain, individual network, and regional (parcel) levels were estimated from resulting adjacency matrices. They included connection strength (median connectivity within and across networks), topological efficiency, robustness, stability and fragility, community structure (modularity), regional connectedness, clustering and centrality (importance of a region in network).

Networks analyzed in this study included large resting-state networks delineated by Yeo et al, which include visual somatomotor, frontopariental control, default, limbic, temporoparietal, dorsal attention and salience networks [73], the thalamus, basal ganglia, and amygdala networks, the cerebellar network, and additional ones including the reward [74], social [75] and central executive networks [76].

#### 2.3.2. Information flow

In addition to estimating non-directional resting-state networks, which are based solely on the strength of connections between regions, we also investigated effective connectivity, i.e., directional interregional connections, measured with phase transfer entropy (PTE) [77], in order to quantify how information is transferred (flows) between regions and is impacted by BMI. Net flow, the difference between afferent and efferent information in a region was also estimated. This analysis was conducted at the resolution of 100 regions, with 3 frames (2.4 s) lag. Pairwise PTE values were computed for all regions, resulting in an asymmetric matrix. For each region, median inflow and outflow were estimated by taking the median over the PTE matrix’s columns and rows, respectively. Directed PTE (dPTE) was also estimated as a normalized measure of directionality, with values >0.5 reflecting higher outflow from the region, and <0.5 higher inflow [78].

#### 2.3.3 Network controllability

Controllability of regional resting-state dynamics was measured by its associated cost, which was estimated using a closed-loop sparsity promoting framework. The latter assumed that resting-state dynamics are controlled by a sparse set of regions that exert their action on the brain even at very high control costs. The framework and estimation of these costs is described in detail in our prior work [66]. Control costs were estimated from time-compressed and thresholded connectivity matrices, assuming that the underlying dynamics can be described by a linear system. Only high control costs associated with a region’s ability to maintain its action over the network were analyzed in the present study. They were assumed to reflect fundamental endogenous mechanisms that regulate brain dynamics and may be aberrantly modulated by excess weight.

#### 2.3.5 Structural MRI data

Morphometric properties included cortical thickness, white matter intensity, and cortical and subcortical volume. They were estimated by the ABCD from fully preprocessed structural MRI (following rigorous quality control), using the gyrus-based Desikan-Killiany cortical atlas [79] to parcellate the brain into 68 regions.

### 2.4 Statistical analysis

Analyses were conducted using generalized linear/logistic regression models with adjustments for age, sex (boys (1) and girls (2)), race (categorized as white, black, mixed race, or other race, given limited statistical power of smaller racial groups, and represented by indicator variables), ethnicity (Hispanic = 1, Non-Hispanic = 0), pubertal stage (represented by an ordinal variable, from pre-puberty (1) to late/post-puberty (4)), family income (provided by the ABCD study as an ordinal variable), physical activity (days per week with ≥60 minutes of exercise), binned sleep duration (from the Sleep Disturbance Scale for Children), and total weekly screen time (other than for homework). In the ABCD study, pubertal stage is calculated based on the Pubertal Development Scale (1-5) [80]. The scale is based on parent’s responses to questions on their child’s physical changes (for example hair growth, breast development, height changes), menarche in females, and deepening voice in males. Given that only few youth were in post-puberty, in statistical models stages 4 and 5 were combined into a single category (= 4). Screen time was calculated as the total weekly number of minutes spent on screens for any purpose other homework and was estimated from the Screen Time Survey. Physical activity was extracted from the Youth Risk Behavior survey.

Analyses were also adjusted for sampling differences across the 21 ABCD sites using study-provided propensity scores. Models with fMRI measures also included adjustments for time of scan acquisition [81] and percent of frames censored for motion in each brain and run.

The significance level was set at = 0.05. All p-values were corrected for the False Discovery Rate [82]. In brain-wide and network-specific analyses, p-values were adjusted for FDR over topological properties (separately for each network). In regional (node-level) analyses, p-values were adjusted for FDR over all nodes in a particular network. Model performance was measured by predictive power. The cohort was randomly partitioned into training and testing sets (75:25; 100 repetitions). The Coefficient of Variation of the Root Mean Square Error (CV[RMSE]) between measurements and predictions was estimated at each repetition. Median (over repetitions) CV[RMSE] was used to assess predictive power. All reported results are based on models with CV[RMSE] ≤0.2. In addition, to maximize reproducibility, at the connectome and network levels, reported results were consistent across two fMRI runs.

However, in some regional analyses, results were only available for the top-quality run, given the computational costs of those analysis. Finally, reported primary results involving continuous BMI/BRI measures were consistent across all these measures, thus ensuring that these results are not BMI measure-dependent.

## 2. RESULTS

Participant demographics are provided in Table 1. Their distribution reflects that the larger ABCD study cohort, which is balanced in terms of sex, but predominantly white and non-Hispanic. Almost 40% of participants were in pre- or early puberty (n = 1,286 (38.5%), 1,091 (32.7%) in mid-puberty, and 785 (23.5%) in late or post-puberty. On average, they were physically active (for ≥60 min) 4.0 days per week (IQR = 3.0), and slept 8-9 hours (IQR = 2 h).

**Table 1.**
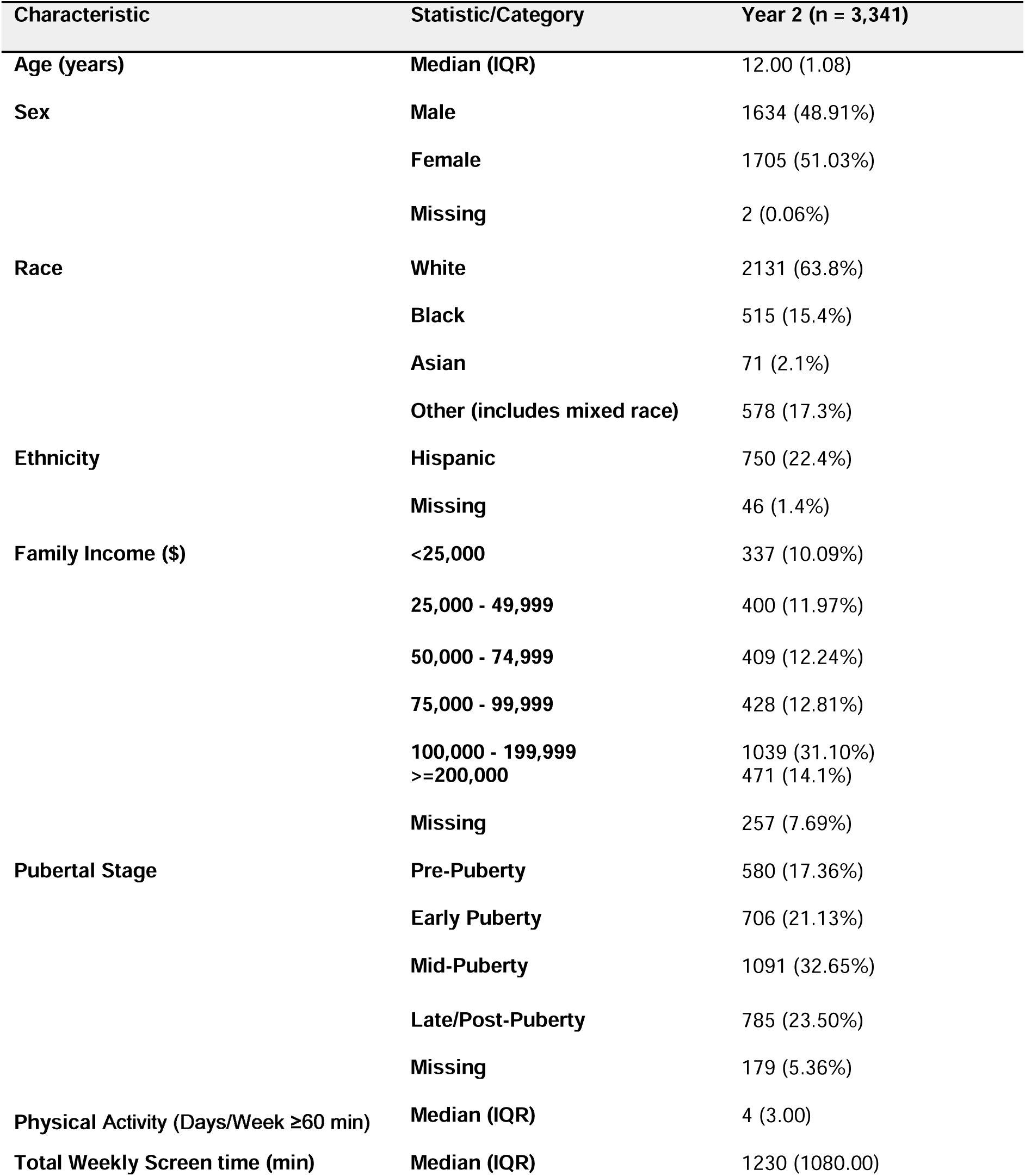
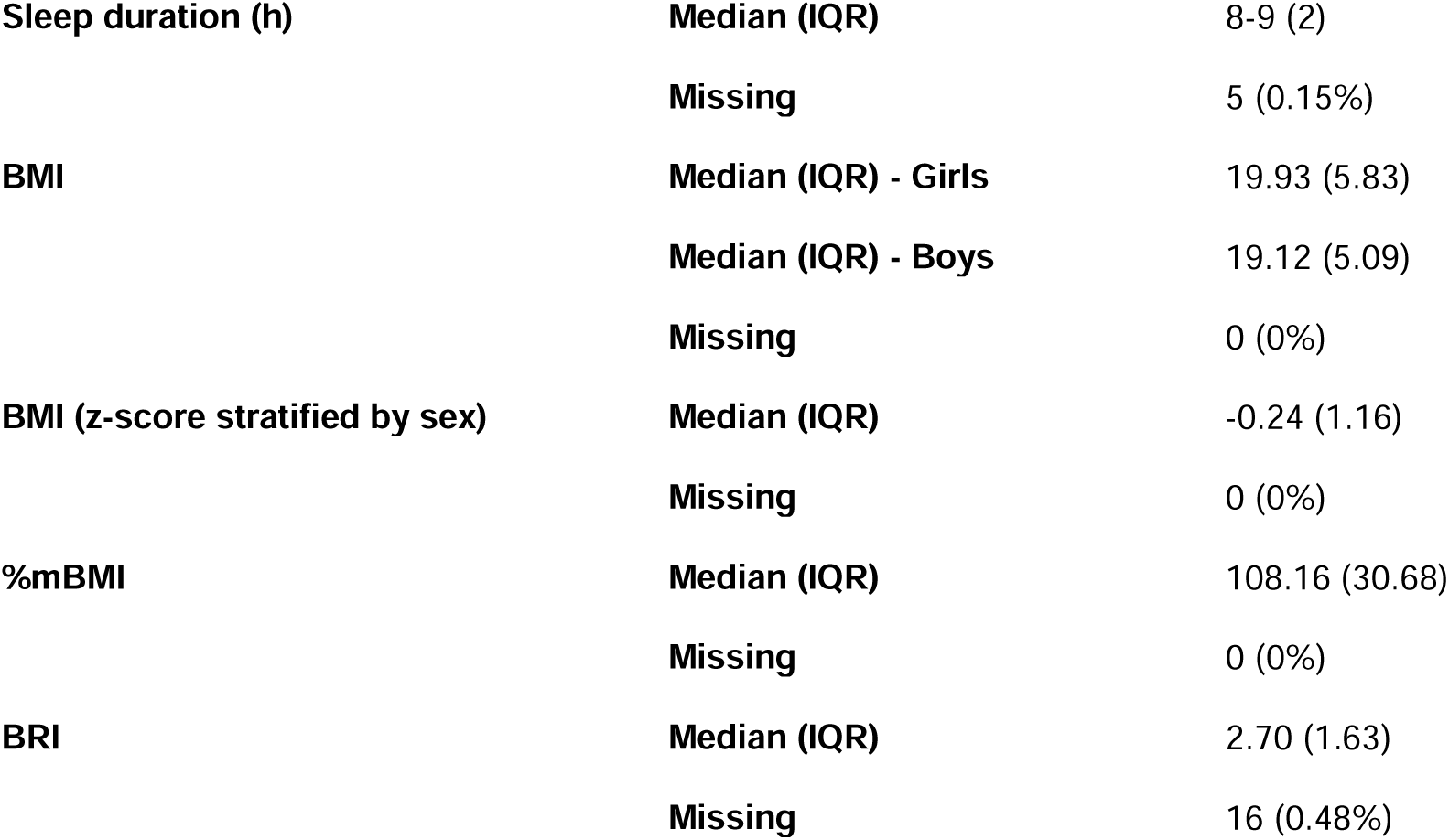
Participant characteristics (n = 3,341). The “Other Non-Hispanic” racioethnic category combines small racial categories: Filipino, Vietnamese, Alaska Native, American Indian, Asian Indian, Chinese, Guamanian, Hawaiian, Japanese, Korean, Native Samoan, other Pacific Islander, other Asian, Multirace, Other race.

Median (IQR) BMI was 19.93 (5.83) kg/m^2^ for girls and 19.12 (5.09) kg/m^2^ for boys, and median (IQR) BRI was 2.70 (1.63). Over 60% of participants had normal BMI (n = 2,212 (66.2%)), 535 (16.0%) had overweight and 594 (17.8%) had obesity. One-hundred-ninety-six (10%) had persistent obesity, 152 (7.8%) had persistent overweight, 164 (8.4%) had normal BMI at baseline but excess at follow-up, and 90 (4.6%) had overweight at baseline and obesity at follow-up (Table 1).

Hispanic youth had higher BMI, %mBMI, and BRI (β=0.19 to 4.95, 95%CI=[0.11, 7.05], p<0.01). Girls, white, and black youth, and those from higher-income families all had lower BMI, %mBMI, and/or BRI (β= −5.28 to −0.14, 95%CI=[−7.22, −0.04], p<0.02),

Higher screen time and shorter sleep were associated with higher BMI, %mBMI, and BRI (β= 0.07, 95%CI=[0.03, 0.11], p<0.01, and β= −0.06, 95%CI=[−0.10, −0.03], p<0.01).

Similar associations were estimated between participant characteristics, overweight, obesity and/or persistent excess BMI (Table 2).

**Table 2.** Statistics of models testing associations between continuous BMI z-score (BMIz), percent of median and participant characteristics. Sex and race-ethnicity statistics are reported as non-standardized betas. All p-values have been adjusted for the False Discovery Rate. CI: 95% Confidence interval. −: Non-significant.

| Variable | BMlz Beta | BMlz CI* | BMlz P-value | %mBMI Beta | %mBMI CI | %mBMI P-value | BRI Beta | BRI CI | BRI P-value |
| --- | --- | --- | --- | --- | --- | --- | --- | --- | --- |
| Pubertal stage | 0.218 | [0.171, 0.264] | <0.001 | 0.220 | [0.173, 0.267] | <0.001 | 0.067 | [0.019, 0.115] | 0.006 |
| Age | - | - | - | -0.069 | [-0.105, -0.032] | <0.001 | 0.043 | [0.004, 0.082] | 0.006 |
| Sex (boys = 0, girls = 1) | -0.194 | [-0.276, -0.111] | <0.001 | -4.084 | [-6.256, -1.912] | <0.001 | - | - | - |
| White | -0.203 | [-0.277, -0.129] | <0.001 | -5.276 | [-7.218, -3.335] | <0.001 | -0.254 | [-0.362, -0.146] | <0.001 |
| Black | - | - | - | - | - | - | -0.193 | [-0.351, -0.035] | 0.017 |
| Hispanic | 0.190 | [0.111, 0.270] | <0.001 | 4.954 | [2.862, 7.046] | <0.001 | 0.416 | [0.300, 0.532] | <0.001 |
| Family income | -0.137 | [-0.177, -0.097] | <0.001 | -0.139 | [-0.179, -0.099] | <0.001 | -0.151 | [-0.193, -0.110] | <0.001 |
| Weekly screen time (h) | 0.072 | [0.027, 0.099] | <0.001 | 0.072 | [0.037, 0.107] | <0.001 | 0.070 | [0.034, 0.105] | <0.001 |
| Sleep duration (h) | -0.063 | [-0.099, -0.027] | <0.001 | -0.064 | [-0.100, -0.028] | <0.001 | -0.063 | [-0.100, -0.026] | <0.001 |
| **BMI STATUS** |  |  |  |  |  |  |  |  |  |
| Variable | Over-weight aOR* | Over-weight CI* | Over-weight P-value | Obesity aOR | Obesity CI | Obesity P-value | BMI History Beta | BMI History CI | BMI History P-value |
| Pubertal stage | 1.275 | [1.119, 1.453] | <0.001 | 1.439 | [1.263, 1.640] | <0.001 | 1.332 | [1.147, 1.548] | <0.001 |
| Sex (boys = 1, girls = 2) | - | - | - | 0.750 | [0.581, 0.968] | 0.027 | - | - | - |
| White | - | - | - | 0.490 | [0.385, 0.624] | <0.001 | 0.547 | [0.414, 0.721] | <0.001 |
| Hispanic | 1.447 | [1.125, 1.860] | 0.004 | 1.693 | [1.336, 2.146] | <0.001 | 1.722 | [1.299, 2.284] | <0.001 |
| Family income | 0.940 | [0.892, 0.991] | 0.022 | 0.869 | [0.829, 0.912] | <0.001 | 0.852 | [0.802, 0.904] | <0.001 |
| Weekly screen time (h) | - | - | - | 1.007 | [1.003, 1.012] | 0.002 | 1.009 | [1.003, 1.015] | 0.003 |
| Sleep duration (h) | - | - | - | 0.836 | [0.741, 0.943] | 0.003 | - | - | - |

### 3.1 Topological associations

#### 3.1.1 Brain-wide and network-specific topology

Higher BMI, %mBMI, BRI, and having obesity were associated with lower brain-wide topological clustering, i.e., lower modular organization of brain circuits (β= −0.07 to −0.05, 95%CI=[−0.11, −0.01], p≤0.03) (Table S1), lower median connectivity (within-and/or cross-network) and/or lower clustering of bilateral frontoparietal control, DM, central executive, and visual networks, and right dorsal attention, salience, and social networks (β= −0.11 to −0.04, 95%CI=[−0.16, −0.01], p≤0.05) and higher fragility of the left central visual network (β= 0.06 to 0.07, 95%CI=[0.02, 0.13], p≤0.04). These BMI measures were also associated with higher fragility (vulnerability of connections) of the thalamus and bilateral basal ganglia (β= 0.05 to 0.10, 95%CI=[0.01, 0.14], p ≤ 0.02), and lower median within-network connectivity of the left basal ganglia (β= −0.05, 95%CI=[−0.09, −0.01], p≤0.02).

Youth with overweight had a more topologically fragile left central visual network (β=0.01, 95%CI=[0.004, 0.013], p<0.01), compared to those with normal BMI, and so did those with persistent excess BMI, i.e., at both assessments (β= 0.01, 95%CI=[0.003, 0.013], p≤0.03). Youth with obesity had lower clustering and/or median within-network connectivity in bilateral central visual and frontoparietal control networks, right peripheral visual, somatomotor, DM, and right social networks, and left central executive and basal ganglia networks (β= −0.015 to −0.002, 95%CI=[−0.024, −0.003], p≤0.03). They also had a more topologically fragile thalamus, bilateral basal ganglia, and the right amygdala-thalamic circuit (β= 0.02 to 0.03, 95%CI=[0.01, 0.05], p≤0.01).

Compared to youth with overweight, those with obesity had lower clustering in the left dorsal attention network and lower median within-network connectivity in the right peripheral visual network (β= −0.01, 95%CI=[−0.02, −0.002], p≤0.04), and more topologically fragile right basal ganglia (β= 0.02, 95%CI=[0.01, 0.04], p≤0.02). Networks with topological properties that were negatively associated with excess BMI, BRI (and/or BMI status) are shown in Figure 1. Model statistics are provided in Table 3, and S2 (associations that were not consistent across all continuous BMI/BRI measures).

**Figure 1.**
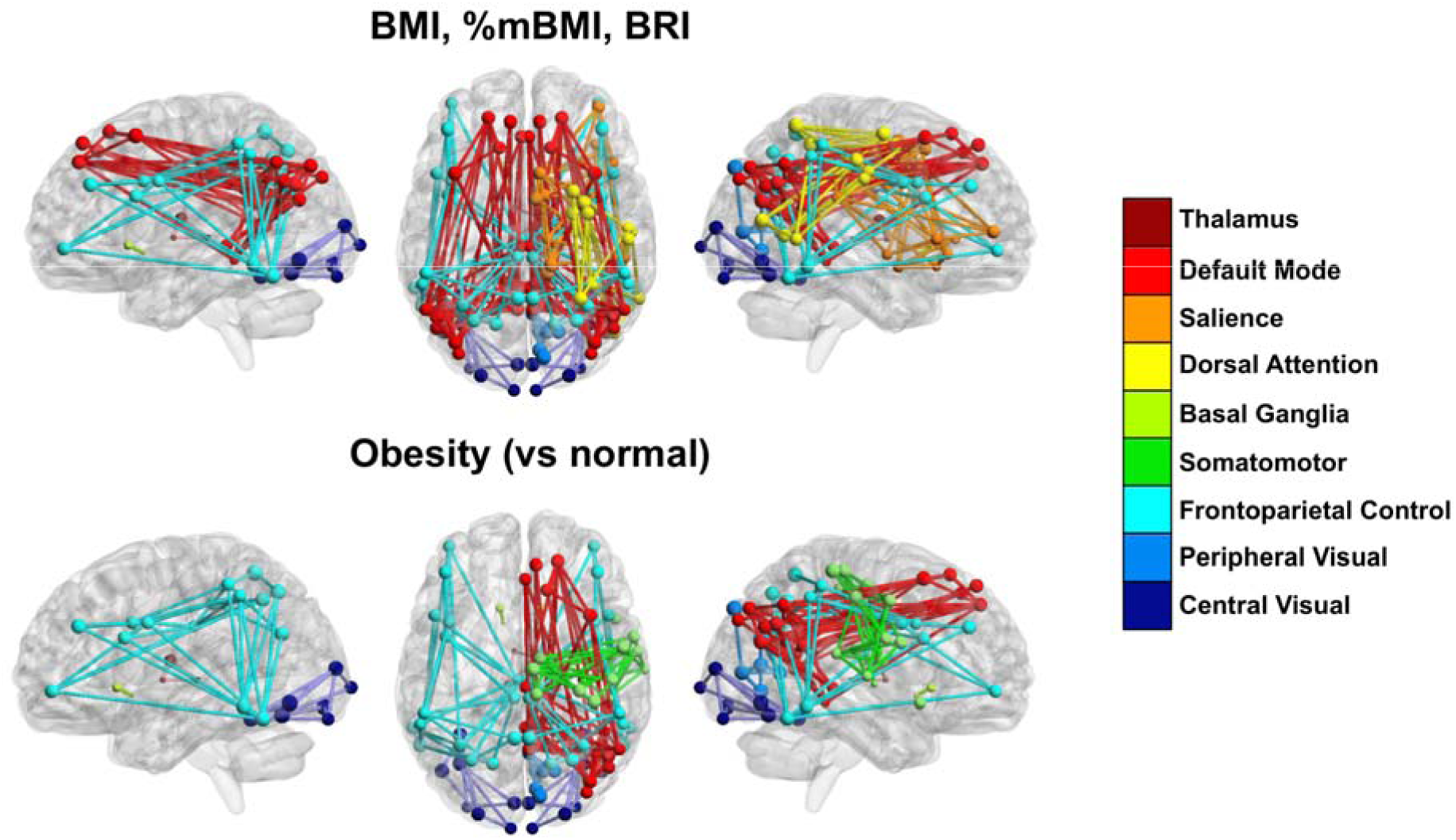
Brain networks with topological properties that are statistically associated with body mass and roundness measures (top) and differ between youth with obesity and those with normal BMI. Each network is shown with a different color. Only a fraction of network connections are shown, and node size corresponds to regional connectedness (degree).

**Table 3.**
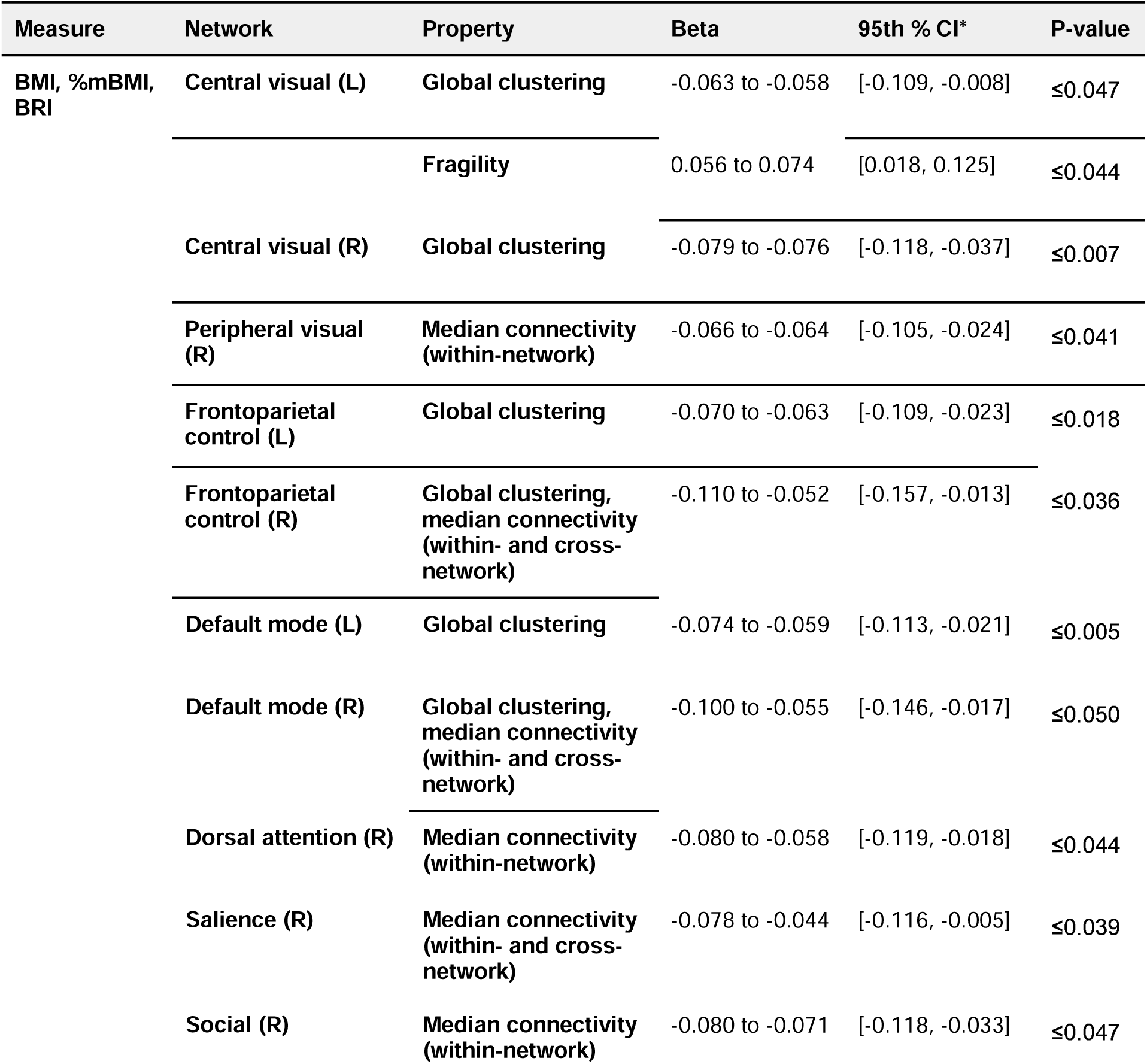
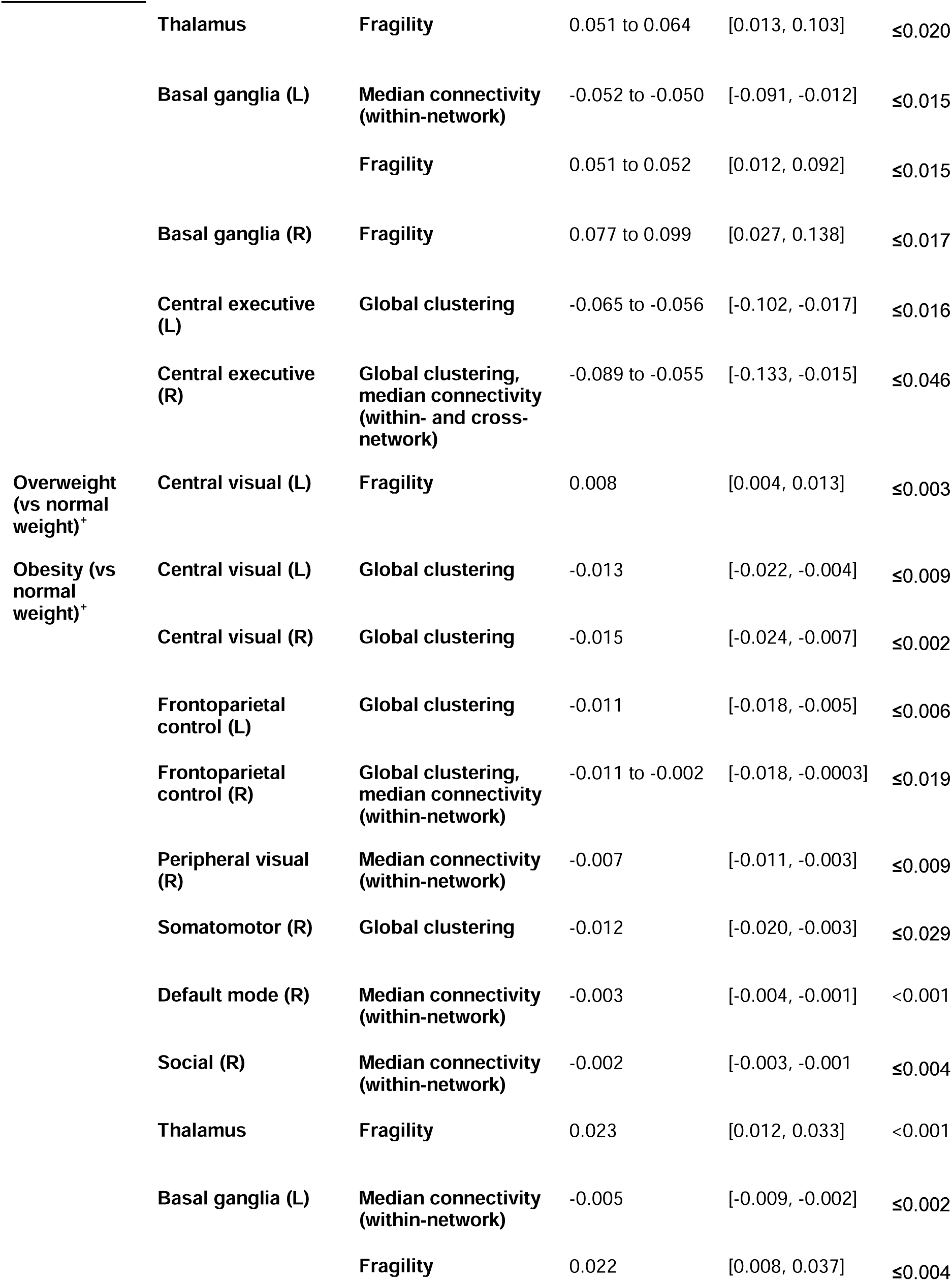
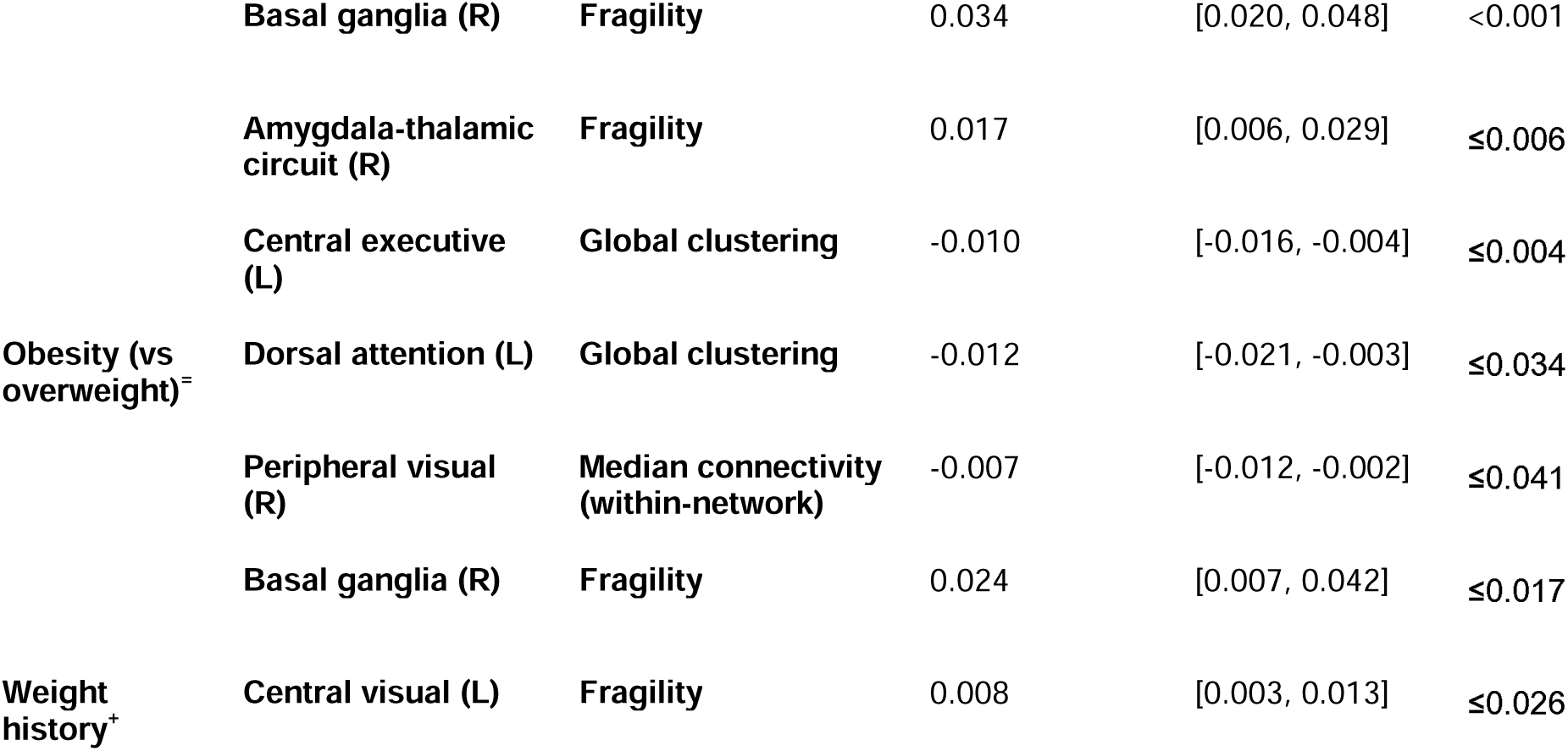
Statistics of multiple linear regression models testing associations between body mass and roundness measures and topological properties of individual networks. All associations were consistent across both runs and had a CV[RMSE] ≤ 0.20. All reported p-values have been adjusted for the False Discovery Rate. ^+^Regression coefficients for binary BMI status variables are not standardized. *-: Nonsignificant; *CI: confidence interval.

#### 3.1.2 Regional topology

Higher BMI, %mBMI, and BRI were associated with lower local connectedness (degree) and/or community structure (local clustering) of regions in the bilateral salience, frontoparietal control, DM, reward, social, prefrontal cortex, reward, and central executive networks. Compared to normal BMI, youth with obesity had lower connectedness and/or clustering in regions of the left dorsal attention, bilateral salience, frontoparietal control, DM, reward, social, prefrontal, and central executive networks, right basal ganglia, and cerebellar networks. The spatial distributions of these associations are shown in Figures 2a (degree) and 2b (clustering).

**Figure 2.**
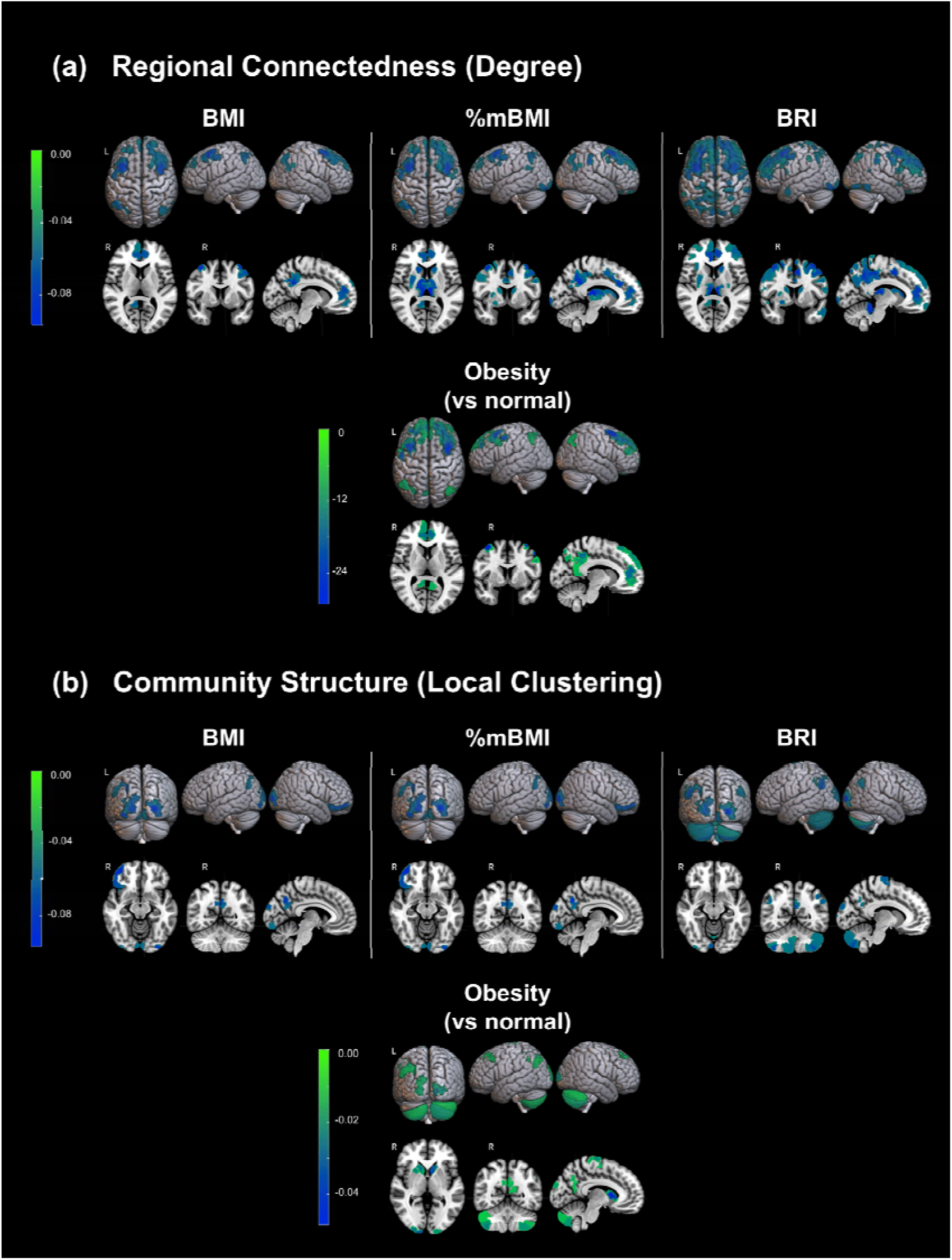
Associations between body mass and roundness measures and (a) regional connectedness (degree − top panels), and (b) community structure (local clustering - bottom panels). The color map represents the values of model regression coefficients; green (highest) to blue (lowest) corresponds to the range of negative coefficients.

### 3.2 Associations with inter-regional brain communication and information transfer

Higher BMI, %mBMI and BRI were associated with median *outflow* from multiple networks and the rest of the brain, including central visual, limbic, frontoparietal control, DM, basal ganglia, amygdala, and thalamus. In comparison to normal BMI, youth with obesity had higher median outflow in regions of central visual, bilateral DM, basal ganglia and thalamus, and right limbic, frontoparietal control, and hippocampus. In comparison to overweight, they had higher median outflow in regions of the right frontoparietal control and right thalamus (Figure 3). Youth with persistent excess BMI had higher median outflow in regions of the bilateral DM network. Corresponding distributions for median *inflow* are shown in Figure S1.

**Figure 3.**
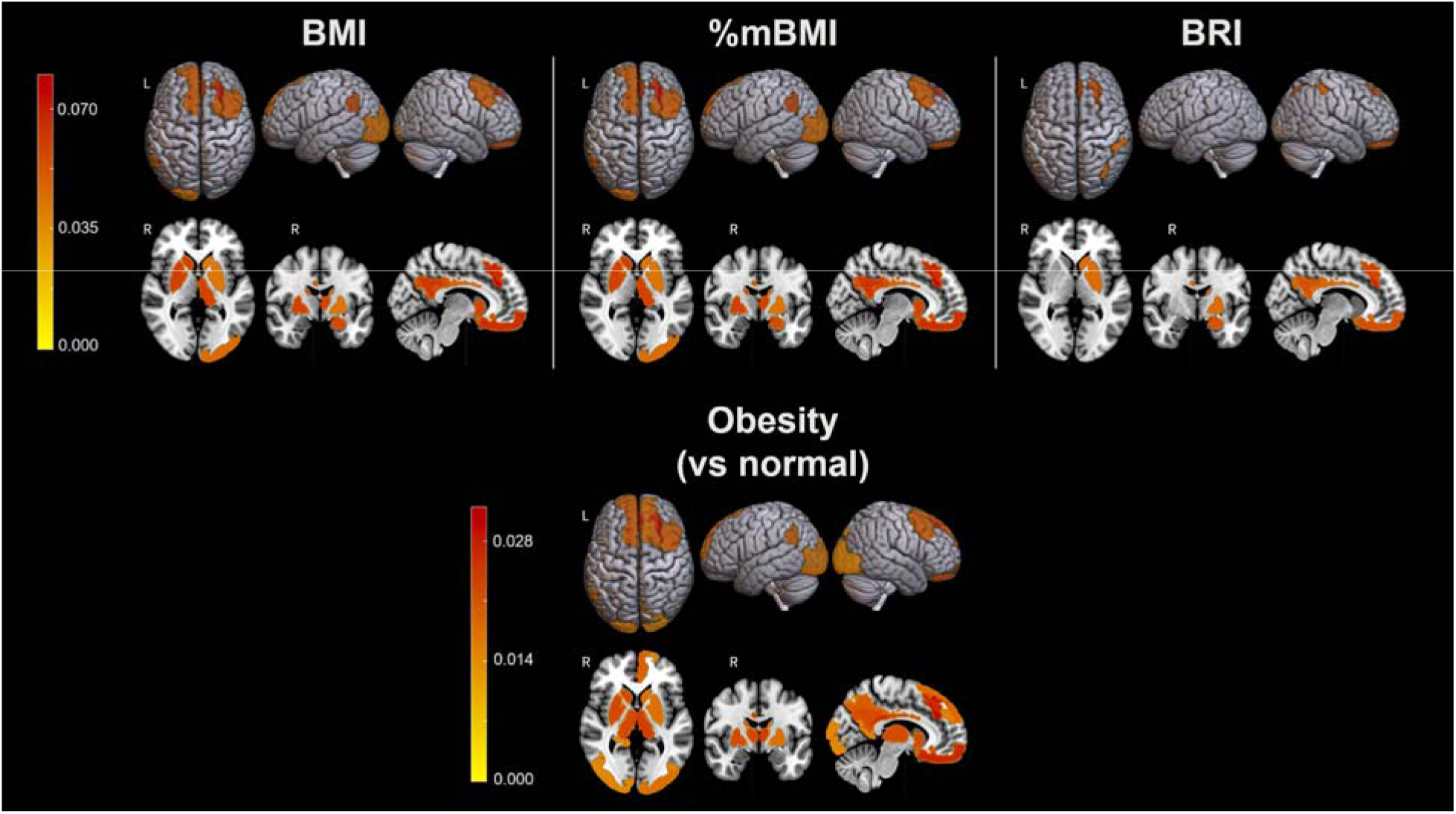
Associations between body mass and roundness measures and median information outflow. The color map represents the values of model regression coefficients; yellow to red corresponds to positive values. Regression coefficients for the binary obesity variable are not standardized.

Higher BMI, %mBMI and BRI were associated with lower overall information flow (measured by median directed transfer entropy (dPTE)) in regions of the right DM network (only BMI and %mBMI) and the cerebellum but higher flow in left central visual, right limbic, and right frontoparietal control regions (Figure 4). Youth with obesity had lower information flow in regions of the salience and somatomotor areas compared to both those with normal BMI and those with overweight. Youth with persistent excess BMI had lower flow in the left cerebellum.

**Figure 4.**
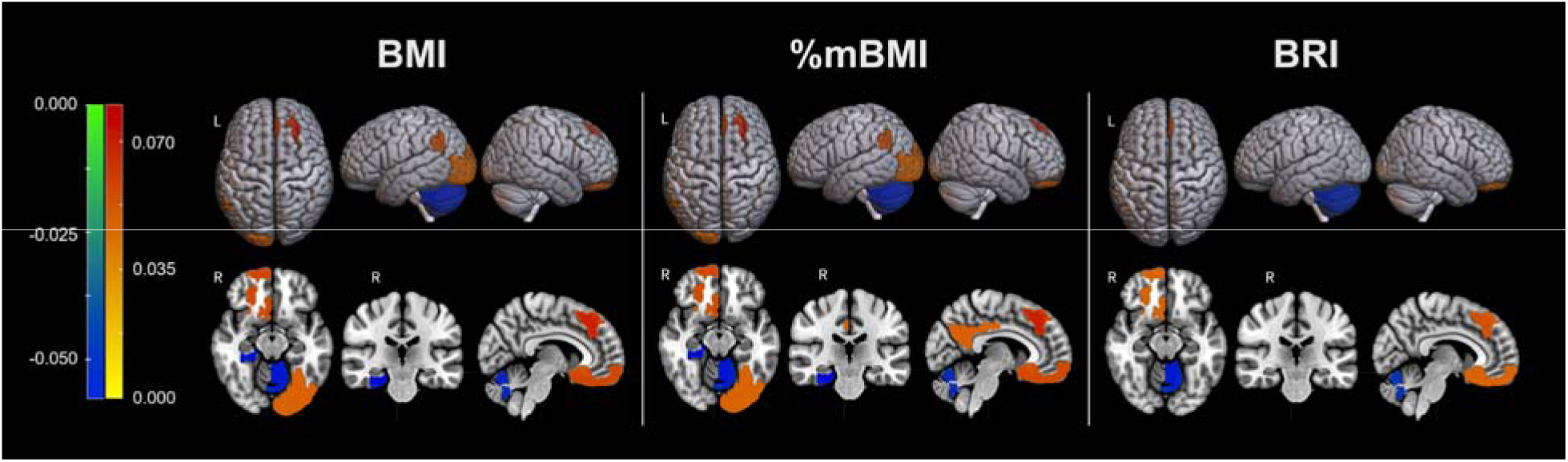
Associations between body mass and roundness measures and overall information flow (measured directed phase transfer entropy (dPTE)). The color map represents the values of model regression coefficients; yellow to red corresponds to positive values and green to blue negative values.

### 3.4 Associations with network controllability

Control costs are associations with a region’s role (and importance) in the controllability of brain dynamics. Youth with higher BMI and BRI had lower control costs associated with regions of left salience and bilateral prefrontal, default-mode, basal ganglia, and thalamic networks, as well as the left precuneus and cingulate (range across measures β = −0.14 to −0.05, 95% CI=[−0.23 to −0.02], p<0.05). The spatial distribution of these associations are shown in Figure 5.

**Figure 5:**
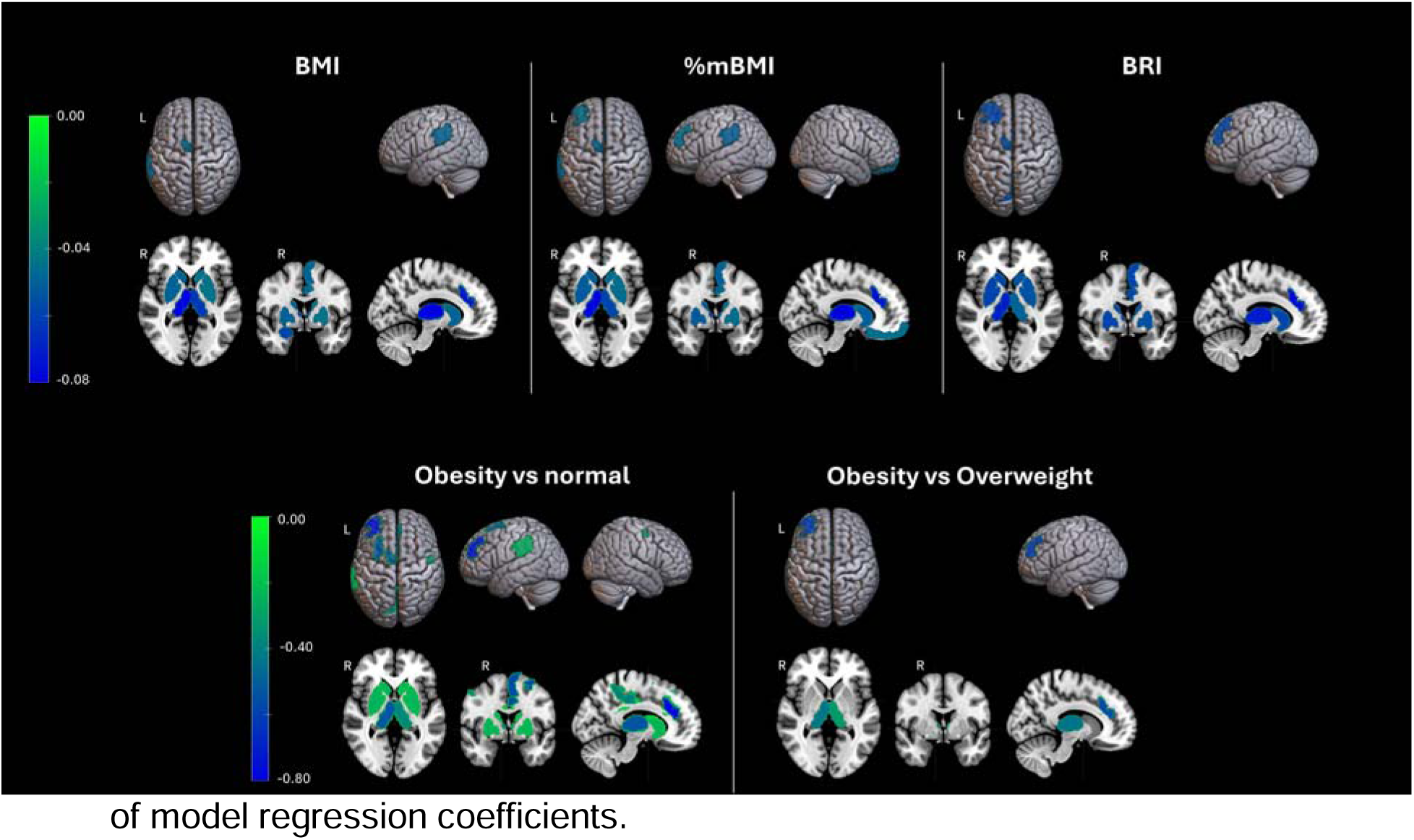
Associations between regional control costs and BMI, %mBMI, BRI, obesity vs normal BMI, and obesity vs overweight. The color map represents the values of model regression coefficients.

### 3.3 Morphometric associations

Higher BMI, %mBMI, and BRI were associated with lower thickness of bilateral pars orbitalis, left paracentral lobule, and right pars triangularis, middle and superior frontal gyrus, and lateral and medial orbitofrontal cortex (β= −0.14 to −0.04, 95%CI=[−0.18, −0.01], p≤0.03); lower volume of the left temporal pole and right rostral middle frontal gyrus (β= −0.09 to −0.04, 95%CI=[−0.13, −0.004], p≤0.04), higher volume of the left cerebellum (β= 0.04 to 0.08, 95%CI=[0.01, 0.12], p≤0.03), lower white matter intensity of the bilateral lateral occipital gyrus (β= −0.11 to −0.06, 95%CI=[−0.15, −0.02], p≤0.02), and higher white matter intensity of the right frontal pole (β= 0.08 to 0.09, 95%CI=[0.04, 0.13], p<0.001). Compared to those with normal BMI, youth with overweight had lower white matter intensity of the left lateral occipital gyrus (β= −0.20, 95%CI=[−0.35, −0.04], p≤0.04), and those with obesity had lower thickness of the bilateral cuneus, paracentral gyrus, frontal pole, and lateral orbitofrontal cortex, right medial orbitofrontal cortex, pars orbitalis, pars triangularis, rostral middle frontal gyrus, and superior frontal gyrus, and left temporal pole (β= −0.04 to −0.02, 95%CI=[−0.06, −0.002], p≤0.05); lower volume of the right superior temporal gyrus, pars triangularis, and rostral middle frontal gyrus, and left amygdala, paracentral gyrus, and temporal pole (β= −341.73 to −26.49, 95%CI=[−608.39, –3.11], p ≤ 0.05), and higher volume of the left cerebellum white matter (β= 296.62, 95%CI=[107.08, 486.17], p<0.01). They also had lower white matter intensity in the left temporal pole, entorhinal cortex, and lateral occipital gyrus (β= −0.36 to −0.29, 95%CI=[−0.59, −0.07], p≤0.04), and higher white matter intensity in bilateral frontal poles (β= 0.29 to 0.45, 95%CI=[0.06, 0.70], p≤0.04). Compared to those with overweight, youth with obesity had lower thickness of the lateral and medial orbitofrontal cortex, paracentral gyrus, pars triangularis, and rostral middle frontal gyrus (β= −0.03 to −0.02, 95%CI=[−0.04, −0.01], p≤0.02), lower volume of the right paracentral gyrus (β= −106.09, 95%CI=[−191.88, −20.30], p=0.02), and higher white matter intensity of the right frontal pole (β= 0.37, 95%CI=[0.04, 0.70], p=0.04).

Youth with persistent excess BMI had lower thickness of the left frontal pole, left temporal pole, right lateral and medial orbitofrontal cortex, right pars orbitalis, right pars triangularis, and right rostral middle frontal gyrus (β= −0.04 to −0.02, 95%CI=[−0.06, − 0.003], p≤0.04), higher thickness of the right postcentral gyrus (β= 0.02, 95%CI=[0.004, 0.04], p=0.04), higher volume of the left isthmus of cingulate gyrus, left cerebellum white matter, left putamen, right thalamus, and right hippocampus (β= 45.53 to 334.99, 95%CI=[0.48, 547.79], p≤0.05); lower white matter intensity of the left lateral occipital gyrus (β= −0.25, 95%CI=[−0.42, −0.08], p=0.01), and higher white matter intensity of the bilateral frontal pole, and right medial orbitofrontal cortex, transverse temporal gyrus, and insula (β= 0.22 to 0.46, 95%CI=[0.02, 0.74], p≤0.05). Model statistics are provided in Table 4 (and measure-specific results in Table S3).

**Table 4.**
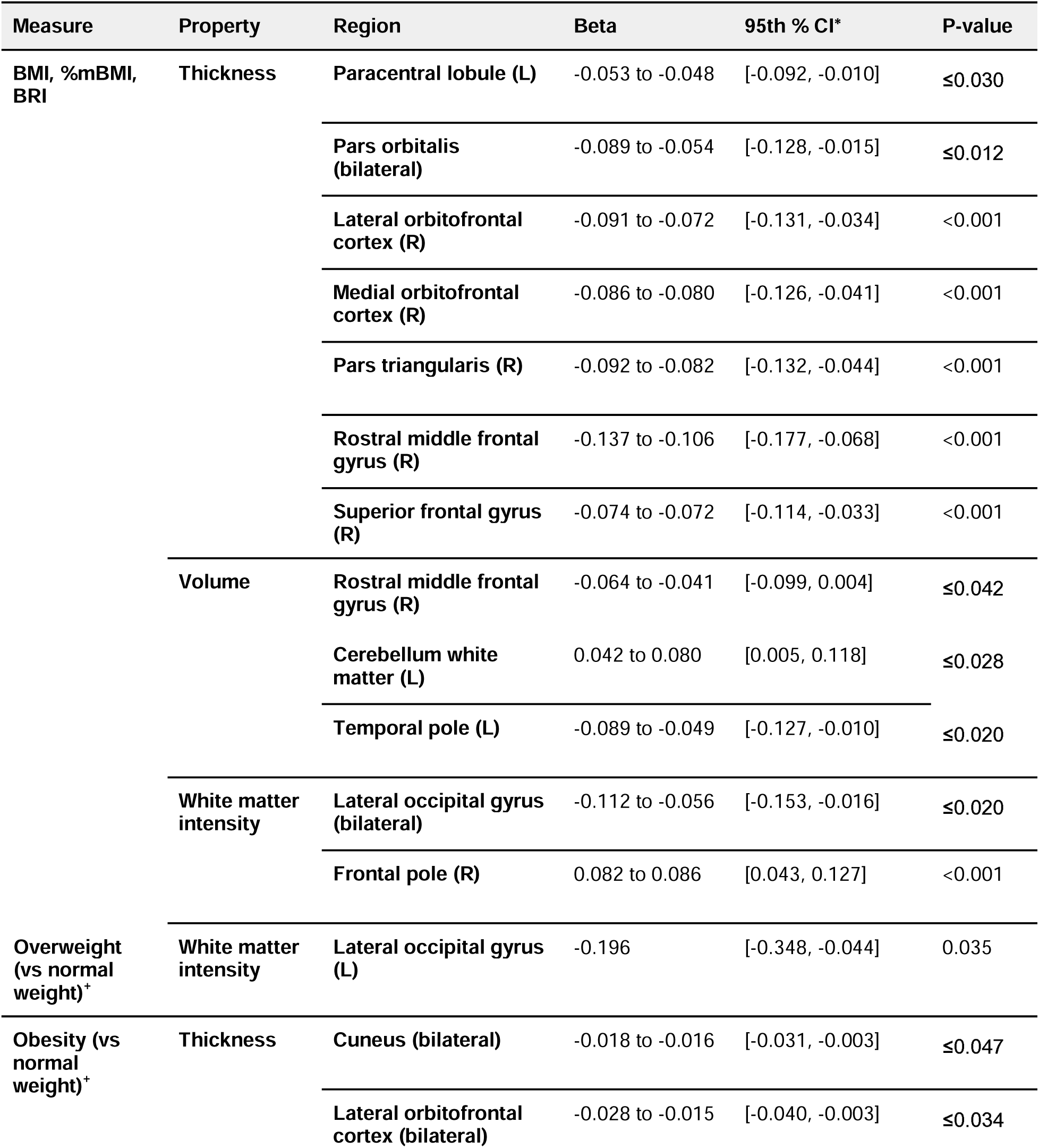
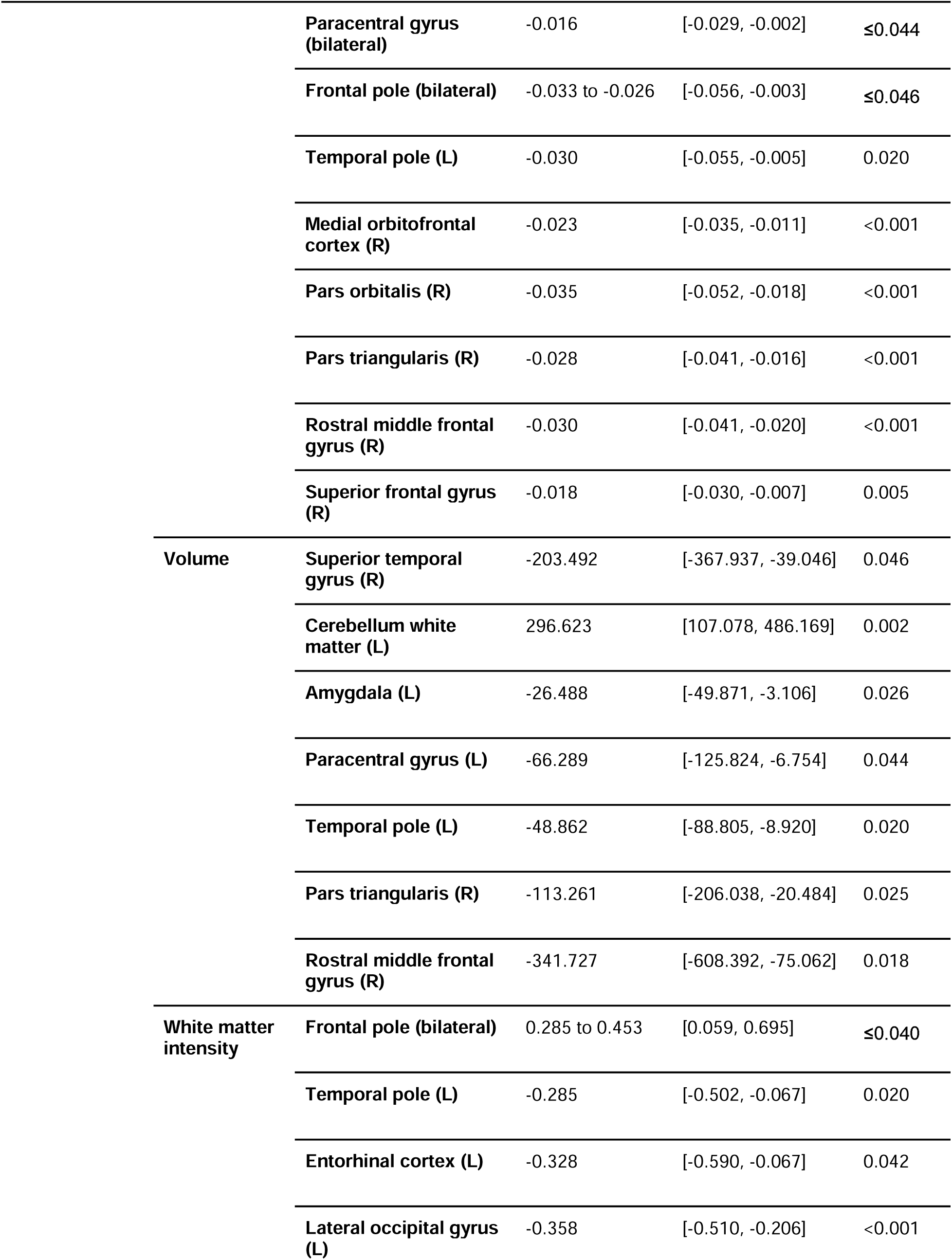
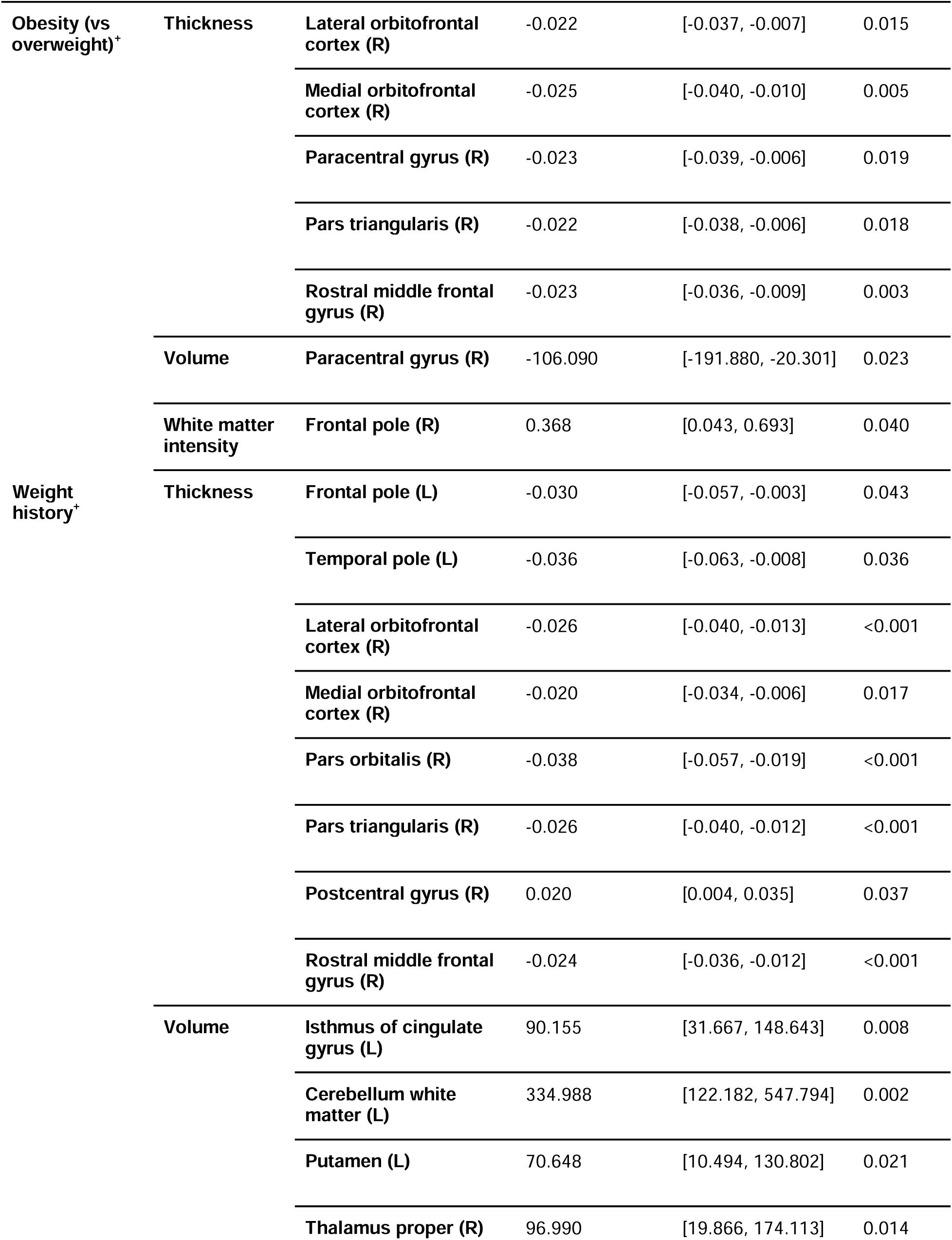
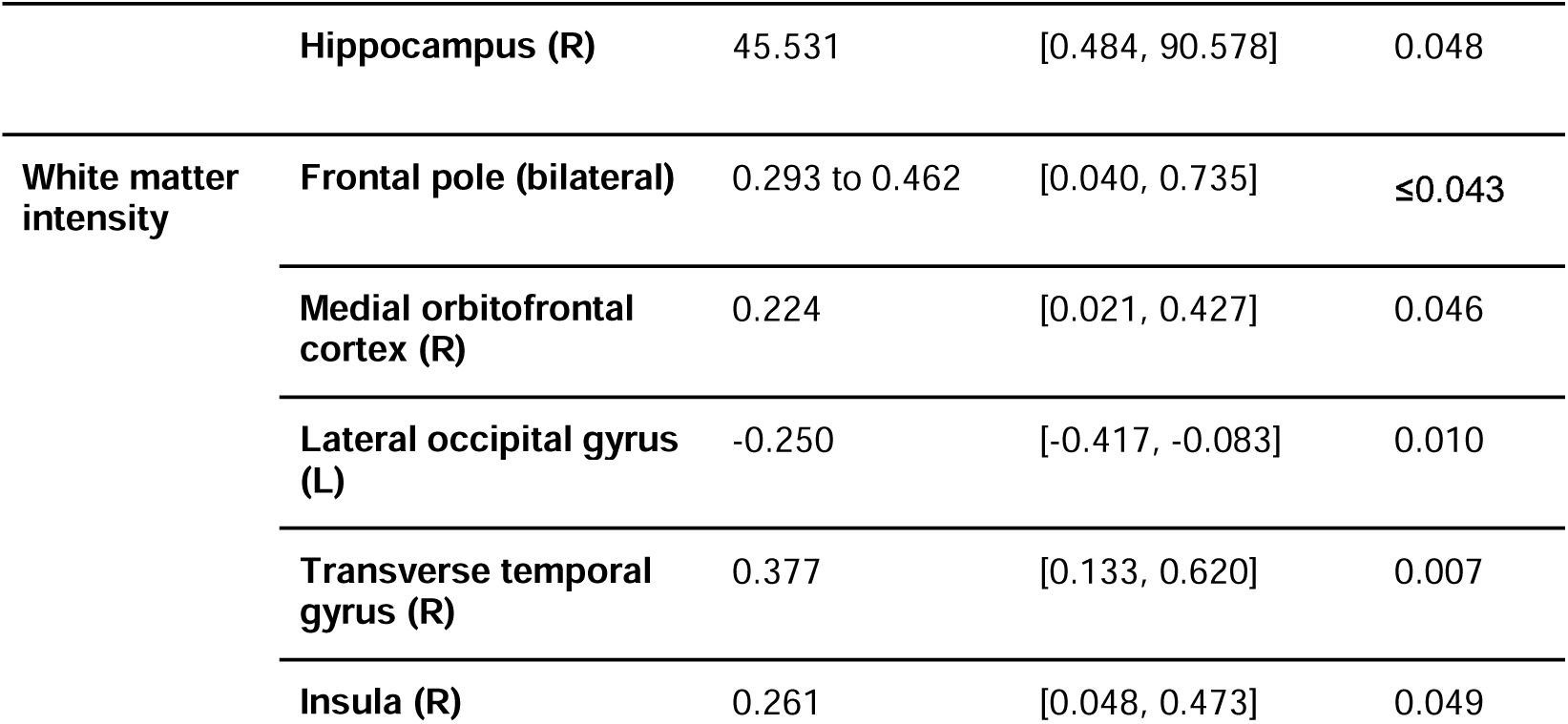
Statistics of multiple linear regression models testing associations between excess weight and morphometric properties. All associations had a CV[RMSE] ≤ 0.20. All reported p-values have been adjusted for the False Discovery Rate. *-: Nonsignificant; *CI: confidence interval. ^+^Regression coefficients for binary BMI status variables are not standardized.

### 3.4 Mappings between functional and structural findings

Lower connectivity and global clustering in the right central executive network were associated with higher cortical thickness of the right pars triangularis and right superior frontal gyrus, respectively (β= −0.06 to −0.04, 95%CI=[−0.10, −0.01], p<0.05). These regions are functionally connected to each other and their properties were negatively associated with BMI, %mBMI, and BRI. Higher cortical thickness of the right pars triangularis was further associated with lower median connectivity in the right frontoparietal control network (β= −0.06, 95%CI=[−0.10, −0.02], p=0.02), and both regions and their properties were negatively associated with BMI, %mBMI, and BRI.

## 3. Discussion

In a large sample of over 3,300 adolescents, our study has addressed an important gap in knowledge, and has investigated associations between excess weight (both present and persistent across 2 years) in adolescents, and the organization of developing resting-state networks (the backbone of the brain’s circuitry), morphological features of their constituent regions, communication between them, and their role in the controllability of its intrinsic dynamics. The study has extended our prior work on the baseline ABCD cohort but has ocused on the two-year follow-up cohort of the ABCD study, has demonstrated consistency of brain-weight associations across BMI and BRI measures (with BRI being a relatively newer but potentially more appropriate measure of adiposity [83]), and has provides important novel insights into detrimental effects of excess weight on structural and topological characteristics of widespread brain networks and regions, interregional communication and information processing, and brain hubs that play critical roles in regulating intrinsic network dynamics and cognitive function.

Youth who spent more time on electronic devices, slept less than recommended, came from lower-income families, or were Hispanic were more likely to have higher BMI and BRI, in agreement with prior studies, including those based on ABCD cohort [84–85].

Higher BMI and BRI, having obesity or (to a lesser extent) overweight were associated with widespread alterations in the organization and inter-regional communication of the frontoparietal control, prefrontal, DM, central executive, salience, dorsal attention and social networks, which undergo significant reorganization in adolescence. These findings have negative implications for cognitive processing across developing domains, including social and executive function, in agreement with prior studies [86–87], and our prior work on the baseline sample of the ABCD [45]. Youth with persistent excess BMI at both baseline and 2-year follow-up had topological alterations (especially increased fragility) in some of the same networks, compared to those with normal BMI across assessments. Topological fragility implies that a circuit is vulnerable to aberrant alterations and impaired connections. Our findings suggest that excess weight may disrupt the maturation of critical brain networks, and increase their vulnerability to stressors.

Another important finding is the association between excess weight/roundness with higher topological fragility and lower strength of regional connections of the thalamus, and the amygdalo-thalamic circuit. The thalamus plays a critical role in cognitive function, but is also specifically involved in feeding, eating behaviors, emotional regulation and reward (including food-related) processing. Impaired connections between the thalamus and cortical and subcortical structures may have detrimental effects on these processes, leading to dysregulation of emotional and reward processing and maladaptive eating behaviors [88]. In addition, higher BMI and BRI were also associated with higher fragility and weaker connections of the basal ganglia (including striatal) networks. Prior studies have reported structural and functional alterations in the thalamus and basal ganglia of youth with obesity, partly resulting from neuroinflammation, that were associated with impaired rewards processing [41][86][89–90]. Our findings suggest complex relationships between excess weight and functional interactions between cortical structures, the thalamus, amygdala and the basal ganglia, which support reward processing, emotional regulation and executive function and inhibitory control [90]. In addition to their topological correlates, high BMI and BRI were associated aberrantly lower control costs in the thalamus and the basal ganglia, indicating that underlying metabolic and/or inflammatory mechanism may modulate fundamental aspects of brain function, such as controllability of its intrinsic dynamics.

Lower control costs were also estimated in regions of the salience, prefrontal, default-mode networks as well as the precuneus and cingulate cortex. Our prior work has shown that some of these regions, including the prefrontal cortex and posterior regions of the default-mode network, maintain their control action on brain network dynamics even at high feedback costs, and overlap with topological and cognitive hubs that play critical roles in the integration of information during cognitive processing [66]. Thus, our findings suggest that excess weight may adversely impact brain hubs that play central regulatory roles in brain function and cognitive processing.

In addition to topological differences, youth with excess BMI also had morphometric differences (including lower volume and thickness), especially in the developing frontal gyrus, medial and/or lateral orbitofrontal cortex, frontal pole, cingulate cortex, insula, pars orbitalis and pars triangularis, in agreement with prior studies [35][91]. These regions undergo significant maturation in adolescence and play critical roles in inhibitory control, decision-making, reward processing and emotional regulation, which are adversely impacted by obesity in youth [36]. A recent study on the ABCD cohort, has identified associations between structural characteristics of the orbitofrontal cortex and food choices and related decision-making [92]. Excess weight was also associated with lower volume, thickness and/or white matter intensity in relatively developed (compared to frontal regions) occipital and sensorimotor areas. This suggests, that adverse metabolic and inflammatory correlates of excess weight may even adversely impact even developed brain areas, with detrimental consequences for sensory processing and sensorimotor integration. Finally, a hallmark characteristic of brain maturation is age-related cortical thinning [93]. We estimated lower cortical thickness in youth with excess BMI in multiple developing brain areas, which suggests potentially accelerated maturation of these areas. However, aberrantly lower cortical thickness has been associated with impaired executive function and mental health issues [94].

Similarly to those with excess weight at the two-year follow-up, those with excess weight at both assessments had structural alterations, including in hub regions, such as the postcentral and cingulate gyri, and the insula. These regions are sparse but extensively connected to the rest of the brain and play central roles in the integration of multidomain information in response to cognitive demands [95–96]. Adverse effects of persistent excess BMI on their morphology may have potentially profound implications for cognitive function across domains, from motor behaviors to emotional regulation and decision-making.

This study has a number of notable strengths, including a large sample, structural and functional neuroimaging data, estimation of the brain’s topological organization beyond connection strength, interregional communication, and investigation of weight effects on network controllability, a fundamental mechanism that is driven by endogenous processes that are likely affected in youth with excess weight. Another strength is the analysis of persistent excess BMI (based on a large subsample with BMI data at two assessments), and importantly the robustness of the results (based on predictive models, consistency across measures (BMI and BRI) and multiple fMRI runs (topological results)). Despite these strengths, the study also has some limitations. As a secondary investigation it is inherently limited by the decisions of the ABCD investigators and available information. For example, information on genetic risk factors for adiposity were not readily available, to facilitate causal analyses. Also, associations between BMI/BRI and environmental stressors were not investigated, as these were outside the scope of the study. A future study could investigate these associations, since the ABCD study collects extensive environmental data. Finally, anthropometric data were not available prior to ages 9-10 years. Therefore, our analyses were limited to two years. As additional assessments become available, future studies could incorporate multiple prior weight measurements, to assess cumulative effects of excess weight on brain development.

This study makes a significant contribution to the fields’ incomplete understanding of the neural underpinnings of adiposity-related cognitive deficits in youth. It shows that that excess weight (and not just obesity but also overweight) may adversely impact hallmark brain characteristics (including markers of neural maturation and controllability of network dynamics), and may impair inter-regional communication and information transfer, especially in functional and cognitive hubs where information is integrated in response to cognitive demands. It also Identified developing circuits and brain regions that are particularly vulnerable to the neuromodulatory effects of adiposity, and could become targets for precise interventions to improve long-term cognitive and mental health outcomes in youth with excess weight.

