## Supplemental materials for "Adiposity in Adolescence has Detrimental Effects on Developing Brain Structures and Organization, Communication, and Controllability of Resting-State Networks"

**FIGURES**

**Figure S1.** Associations between body mass and roundness measures and median information inflow. The color map represents the values of model regression coefficients; yellow to red corresponds to positive values. Regression coefficients for the binary obesity variables are not standardized.

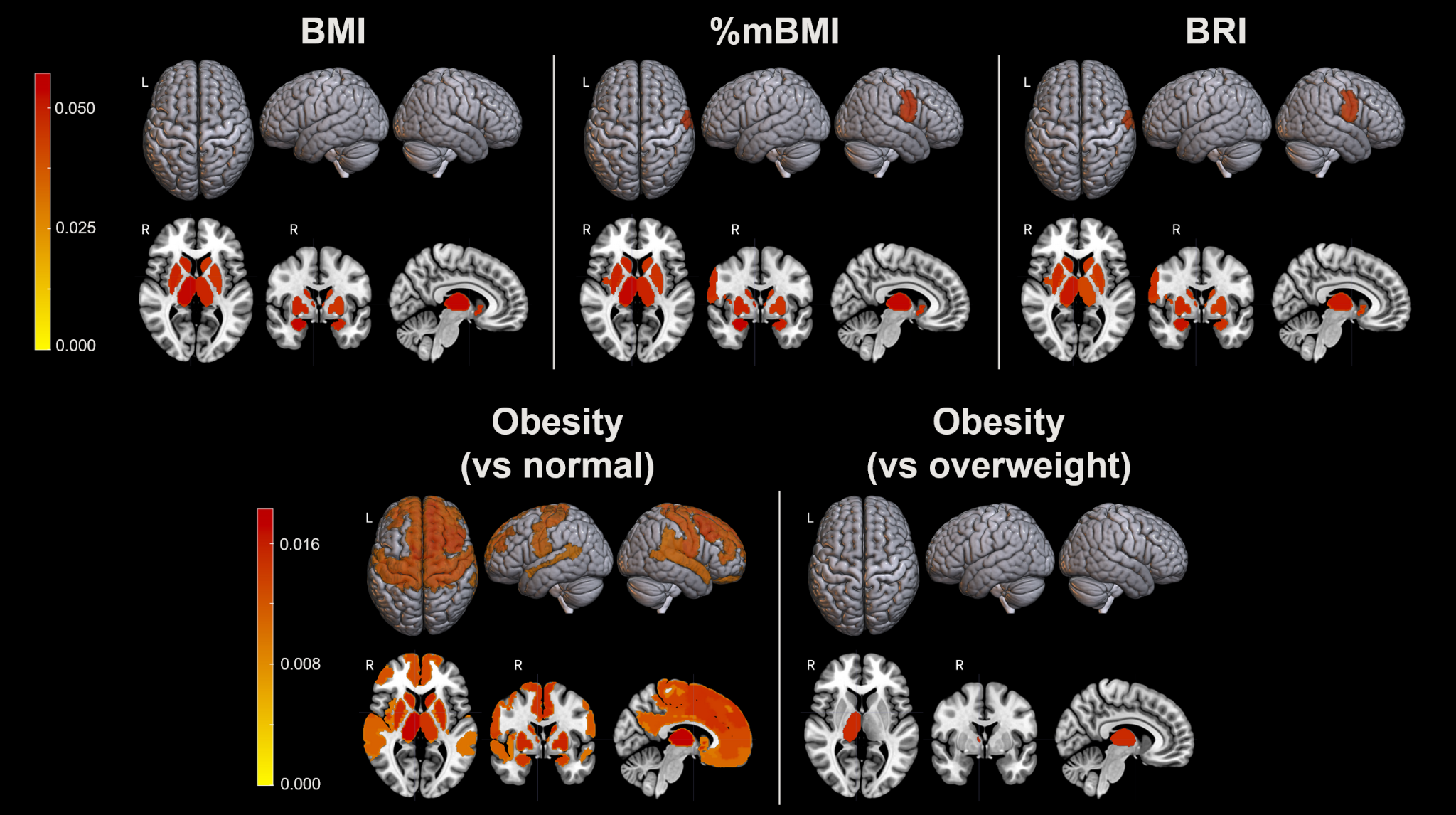

**TABLES**

**Table S1.** Statistics of multiple linear regression models testing associations between body mass and roundness measures and whole-brain topological properties. All associations were consistent across both runs and had a CV[RMSE] ≤ 0.20. All reported p-values have been adjusted for the False Discovery Rate. *-: Nonsignificant; *CI: confidence interval. ^+^Regression coefficients for binary BMI status variables are not standardized.

| **Measure** | **Property** | **Beta** | **95th % CI*** | **P-value** |
| --- | --- | --- | --- | --- |
| **BMI** | **Global clustering** | -0.053 | [-0.092, -0.013] | 0.029 |
| **%mBMI** | **Global clustering** | -0.053 | [-0.092, -0.014] | 0.027 |
| **BRI** | **Global clustering** | -0.070 | [-0.108, -0.032] | 0.002 |
|  | **Median connectivity** | -0.043 | [-0.081, -0.004] | 0.048 |
|  | **Topological robustness** | -0.048 | [-0.086, -0.010] | 0.026 |
|  | **Topological stability** | -0.048 | [-0.086, -0.010] | 0.026 |
| **Obesity (vs normal weight)** | **Global clustering** | -0.006^+^ | [-0.010, -0.002] | 0.018 |

**Table S2.** Statistics of multiple linear regression models testing associations between body mass and roundness measures and topological properties of individual networks which were not consistent across the continuous measures. All associations were consistent across both runs and had a CV[RMSE] ≤ 0.20. All reported p-values have been adjusted for the False Discovery Rate. Regression coefficients for binary BMI status variables are not standardized. *-: Nonsignificant; *CI: confidence interval.

| **Measure** | **Network** | **Property** | **Beta** | **95th % CI*** | **P-value** |
| --- | --- | --- | --- | --- | --- |
| **BMI** | **Default mode (L)** | **Median connectivity (within-network)** | -0.074 to -0.070 | [-0.113, -0.032] | ≤0.002 |
|  | **Amygdala-thalamic circuit (R)** | **Fragility** | 0.057 | [0.016, 0.097] | ≤0.015 |
| **%mBMI** | **Amygdala-thalamic circuit (R)** | **Fragility** | 0.057 | [0.016, 0.097] | ≤0.014 |
| **BRI** | **Central visual (L)** | **Median connectivity (within-network)** | -0.063 to -0.043 | [-0.102, -0.004] | ≤0.044 |
|  | **Peripheral visual (L)** | **Median connectivity (within-network)** | -0.076 | [-0.115, -0.037] | ≤0.001 |
|  | **Somatomotor (L)** | **Median connectivity (within-network)** | -0.061 | [-0.099, -0.023] | ≤0.010 |
|  | **Dorsal attention (L)** | **Global clustering, median connectivity (within-network)** | -0.057 to -0.048 | [-0.096, -0.010] | ≤0.048 |
|  | **Default mode (L)** | **Median connectivity (within-network)** | -0.073 to -0.072 | [-0.112, -0.035] | <0.001 |
|  | **Central visual (R)** | **Median connectivity (within-network)** | -0.076 to -0.053 | [-0.115, -0.014] | ≤0.015 |
|  | **Somatomotor (R)** | **Median connectivity (within- and cross-network)** | -0.060 to -0.049 | [-0.098, -0.011] | ≤0.023 |
|  |  | **Fragility** | 0.055 | [0.017, 0.094] | ≤0.013 |
|  | **Dorsal attention (R)** | **Global clustering, median connectivity (cross-network)** | -0.080 to -0.053 | [-0.119, -0.015] | ≤0.016 |
|  | **Reward (R)** | **Median connectivity (within-network)** | -0.070 | [-0.108, -0.032] | <0.001 |

**Table S3.** Statistics of multiple linear regression models testing associations between body mass and roundness measures and morphometric properties which were not consistent across the continuous measures. All associations had a CV[RMSE] ≤ 0.20. All reported p-values have been adjusted for the False Discovery Rate. *-: Nonsignificant; *CI: confidence interval.

| **Measure** | **Property** | **Region** | **Beta** | **95th % CI*** | **P-value** |
| --- | --- | --- | --- | --- | --- |
| **BMI** | **Thickness** | **Inferior parietal gyrus (L)** | -0.044 | [-0.084, -0.004] | 0.045 |
|  |  | **Lateral orbitofrontal cortex (L)** | -0.061 | [-0.101, -0.022] | 0.008 |
|  |  | **Rostral middle frontal gyrus (L)** | -0.050 | [-0.090, -0.010] | 0.043 |
|  |  | **Frontal pole (L)** | -0.056 | [-0.096, -0.016] | 0.009 |
|  |  | **Temporal pole (L)** | -0.047 | [-0.087, -0.006] | 0.024 |
|  |  | **Insula (L)** | 0.051 | [0.011, 0.091] | 0.037 |
|  |  | **Cuneus (R)** | -0.051 | [-0.090, -0.012] | 0.030 |
|  |  | **Frontal pole (R)** | -0.066 | [-0.106, -0.026] | 0.002 |
|  | **Volume** | **Parahippocampal gyrus (bilateral)** | 0.050 to 0.059 | [0.011, 0.098] | ≤0.040 |
|  |  | **Pallidum (L)** | 0.046 | [0.008, 0.084] | 0.017 |
|  |  | **Cerebellum white matter (R)** | 0.052 | [0.014, 0.090] | 0.008 |
|  |  | **Thalamus proper (R)** | 0.053 | [0.016, 0.089] | 0.005 |
|  | **White matter intensity** | **Inferior parietal gyrus (L)** | -0.050 | [-0.090, -0.009] | 0.045 |
|  |  | **Frontal pole (L)** | 0.062 | [0.022, 0.103] | 0.008 |
|  |  | **Temporal pole (L)** | -0.067 | [-0.107, -0.027] | 0.003 |
|  |  | **Entorhinal cortex (bilateral)** | -0.062 to -0.053 | [-0.102, -0.012] | ≤0.031 |
| **%mBMI** | **Thickness** | **Inferior parietal gyrus (L)** | -0.045 | [-0.085, -0.006] | 0.038 |
|  |  | **Lateral orbitofrontal cortex (L)** | -0.062 | [-0.102, -0.022] | 0.007 |
|  |  | **Rostral middle frontal gyrus (L)** | -0.050 | [-0.089, -0.010] | 0.043 |
|  |  | **Frontal pole (bilateral)** | -0.067 to -0.056 | [-0.107, -0.016] | ≤0.009 |
|  |  | **Temporal pole (L)** | -0.046 | [-0.087, -0.006] | 0.025 |
|  |  | **Insula (L)** | 0.050 | [0.010, 0.090] | 0.041 |
|  |  | **Cuneus (R)** | -0.050 | [-0.089, -0.011] | 0.034 |
|  |  | **Pars opercularis (R)** | -0.049 | [-0.089, -0.009] | 0.048 |
|  | **Volume** | **Parahippocampal gyrus (bilateral)** | 0.049 to 0.059 | [0.010, 0.098] | 0.042 |
|  |  | **Pallidum (L)** | 0.046 | [0.009, 0.084] | 0.016 |
|  |  | **Cerebellum white matter (R)** | 0.052 | [0.014, 0.090] | 0.007 |
|  |  | **Thalamus proper (R)** | 0.053 | [0.016, 0.089] | 0.005 |
|  | **White matter intensity** | **Inferior parietal gyrus (L)** | -0.049 | [-0.089, -0.008] | 0.038 |
|  |  | **Frontal pole (L)** | 0.063 | [0.023, 0.104] | 0.006 |
|  |  | **Temporal pole (L)** | -0.066 | [-0.106, -0.026] | 0.004 |
|  |  | **Entorhinal cortex (bilateral)** | -0.061 to -0.052 | [-0.102, -0.012] | ≤0.035 |
| **BRI** | **Thickness** | **Cuneus (L)** | -0.057 | [-0.095, -0.020] | 0.009 |
|  |  | **Lingual gyrus (L)** | -0.046 | [-0.083, -0.009] | 0.031 |
|  |  | **Paracentral gyrus (R)** | -0.050 | [-0.089, -0.012] | 0.015 |
|  |  | **Posterior cingulate cortex (R)** | -0.048 | [-0.087, -0.009] | 0.047 |
|  | **Volume** | **Lingual gyrus (L)** | -0.044 | [-0.081, -0.007] | 0.031 |
|  |  | **Paracentral gyrus (bilateral)** | -0.049 to -0.047 | [-0.086, -0.009] | ≤0.023 |
|  |  | **Pars orbitalis (L)** | -0.059 | [-0.095, -0.022] | 0.005 |
|  |  | **Rostral middle frontal gyrus (L)** | -0.049 | [-0.084, -0.014] | 0.018 |
|  |  | **Superior frontal gyrus (bilateral)** | -0.054 to -0.053 | [-0.090, -0.018] | ≤0.008 |
|  |  | **Superior temporal gyrus (bilateral)** | -0.075 to -0.064 | [-0.111, -0.028] | ≤0.001 |
|  |  | **Caudal middle frontal gyrus (R)** | -0.055 | [-0.093, -0.018] | 0.012 |
|  |  | **Lateral orbitofrontal cortex (R)** | -0.046 | [-0.081, -0.010] | 0.017 |
|  |  | **Medial orbitofrontal cortex (R)** | -0.038 | [-0.074, -0.002] | 0.038 |
|  |  | **Pars triangularis (R)** | -0.066 | [-0.103, -0.028] | <0.001 |
|  |  | **Temporal pole (R)** | -0.048 | [-0.086, -0.011] | 0.036 |
|  |  | **Amygdala (L)** | -0.054 | [-0.091, -0.018] | 0.004 |
|  | **White matter intensity** | **Lateral orbitofrontal cortex (R)** | 0.043 | [0.004, 0.082] | 0.033 |
|  |  | **Medial orbitofrontal cortex (R)** | 0.057 | [0.018, 0.096] | 0.007 |
|  |  | **Insula (R)** | 0.049 | [0.009, 0.088] | 0.048 |
